# Host-parasite dynamics set the ecological theatre for the evolution of state- and context-dependent dispersal in hosts

**DOI:** 10.1101/2020.04.03.022962

**Authors:** Jhelam N. Deshpande, Oliver Kaltz, Emanuel A. Fronhofer

## Abstract

While host-parasite interactions are ubiquitous, the large scale consequences of parasite infections are mainly driven by the spatial context. One trait of pivotal importance for the eco-evolutionary dynamics of such metapopulations is the spatial behaviour of hosts, that is, their dispersal. It is well established that dispersal is not a random process, rather dispersal is informed and may depend on internal and external factors. In host-parasite metapopulations, dispersal may be a function of a host’s infection state, but also of the local context, such as host density or parasite prevalence. Using a dynamical host-parasite metapopulation model, we explore whether host dispersal evolves to be state- and context-dependent and what shapes the evolutionarily stable dispersal reaction norms have. We show that state-dependent dispersal readily evolves in the sense that hosts disperse more when infected. This dispersal bias evolves due to kin selection which is consistent with previous studies. Most importantly, we show that prevalence-dependent dispersal evolves, especially when virulence is high and epidemiological dynamics have predictable signatures. The observed evolutionary outcome, a negatively prevalence-dependent dispersal reaction norm for susceptible hosts, seems counter-intuitive at first. However, our results can be readily explained by the emergent epidemiological dynamics, especially their spatial and temporal correlation patterns. Finally, we show that context-dependency in dispersal may rely on both, prevalence, but also host density cues. Our work provides new insights into the evolution of complex dispersal phenotypes in host-parasite metapopulations as well as on associated feedbacks between ecological dynamics and evolutionary change.

## Introduction

In many host-parasite systems, spatial behaviour of hosts, more precisely, their movement and dispersal, is key to the spread of infection, especially if populations are spatially structured. In such spatially structured populations (metapopulation *sensu lato*), the landscape is simplified and represented as a network of interconnected patches (nodes), where the probability of local transmission depends, for instance, on the global rates at which infected and uninfected individuals move between patches, the direction they choose, or the distances they cover (e.g. Kerr et al., 2006; Ferrari et al., 2008; Metcalf et al., 2013; Yashima and Sasaki, 2014; Dudas et al., 2017). A metapopulation approach of disease transmission is not only important for understanding and predicting the spread of epidemics across landscapes (Ostfeld et al., 2005; Keeling and Eames, 2005; North and Godfray, 2017). Variation in host and parasite dispersal may also have a strong impact on the evolution of attack and defence traits, such as virulence and resistance, or patterns of local adaptation (reviewed in Lion and Gandon, 2015, 2016; Penczykowski et al., 2016). Typically, however, in the host-parasite literature dispersal is considered as a fixed parameter in theoretical models or set by the experimenter in empirical studies (e.g. Morgan et al., 2005). Therefore, it is largely unknown how dispersal itself may evolve in response to interactions with parasites, and how this might feed back on to epidemiological and coevolutionary processes.

Dispersal has long been recognised as an important driver of metapopulation dynamics (Hanski, 1999; Clobert et al., 2012), and over recent years there has been an increasing interest in the evolution of dispersal (Bowler and Benton, 2005; Ronce, 2007; Saastamoinen et al., 2018). It is now well established that dispersal has a genetic basis (Saastamoinen et al., 2018). Thus, dispersal may readily evolve in response to spatial variation in fitness expectations (Bowler and Benton, 2005), resulting from habitat variability (Holt and McPeek, 1996), kin competition (Hamilton and May, 1977), or inbreeding depression (Bengtsson, 1978; Pusey and Wolf, 1996; Perrin and Mazalov, 2000). Only recently, theoretical work has begun to address dispersal evolution among interacting species in metacommunities (e.g. Poethke et al., 2010; Chaianunporn and Hovestadt, 2012b,a; Amarasekare, 2016).

In the case of interactions with natural enemies, such as predators, herbivores or parasites, one may expect evolution of both constitutive and plastic dispersal behaviour. For example, constitutive long-distance dispersal in plants may evolve as a means to escape from specialised, locally dispersing pests and herbivores (Muller-Landau et al., 2003). More recently, Chaianunporn and Hovestadt (2012b) showed that, if predation increases prey population extinction risk, higher constitutive dispersal rates may evolve to compensate this risk.

In addition to such constitutive evolutionary changes, host dispersal behaviour may have evolved to be plastic in two main ways (see also Clobert et al., 2009): Dispersal may depend on the internal state of the host (state-dependence), that is, being infected or not, for example. In addition, dispersal may depend on the external context (context-dependence) which includes host densities or parasite prevalences, to name but two examples. Examples for the first case, that is, dispersal being infection state-dependent have been reported for various organisms, from protists to vertebrates (Sorci et al., 1994; Heeb et al., 1999; Suhonen et al., 2010; Fellous et al., 2011; Brown et al., 2016). However, the adaptive significance of these changes is not always clear. Reduced dispersal of infected hosts (e.g. Fellous et al., 2011) may simply be a physiological by-product of exploitation of host resources by the parasite. Conversely, increased dispersal may reflect manipulation by the parasite to increase parasite dispersal (Lion et al., 2006; Kamo and Boots, 2006), rather than evolution of the host. Evolution of host dispersal plasticity has been studied theoretically by Iritani and Iwasa (2014) and Iritani (2015). They find that ‘infection-biased’ dispersal, that is, increased dispersal when infected, is favoured with increasing probability of recovery during dispersal, with decreasing dispersal costs, and with higher levels of parasite virulence. Kin selection is the driving force underlying these predictions because the dispersal of infected individuals reduces the negative effects of kin competition on philopatric susceptibles, similar to Hamilton and May (1977).

In the latter case of dispersal plasticity, that is, context-dependent dispersal of hosts, dispersal occurs in response to the presence of natural enemies in a population, and the selective advantage is the possibility of escaping from mortality, just like in the above models of constitutive dispersal evolution. Examples of context-dependent dispersal are known from plants and animals where the presence of herbivores or predators induces an increased production of long-distance seeds (e.g. de la Pena and Bonte, 2014) or a higher proportion of winged morphs in insects (e.g. Weisser et al., 1999). This type of informed dispersal requires the capacity to ‘sense’ the presence of the enemy, either through direct contact when leaves are eaten by herbivores, for example, or indirect queues which can be mediated by chemicals released by predators or parasites or by damaged and infected hosts, for instance. Fronhofer et al. (2018) showed that such chemical predator-related cues can indeed induce increased dispersal in various invertebrate and vertebrate organisms.

Despite the above discussed empirical examples, we are not aware of theoretical treatments of the evolution of context-dependent dispersal in host-parasite systems. Results of predator-prey models indicate that conditions for escape dispersal to evolve may be restrictive (Poethke et al., 2010; Hammill et al., 2015). Namely, emigration from a currently infected population may not have a selective advantage if the probability of encountering the parasite is similar in other populations, or if encounter probabilities are unpredictable altogether. Moreover, if infected individuals take the same dispersal decisions as uninfected ones, escape benefits may be further limited.

Here, we develop theory for the joint evolution of state-dependent and context-dependent dispersal of hosts facing infectious disease. Using a spatially explicit, individual-based host-parasite model, we investigate conditions for the evolution of infection-state dependent dispersal and prevalence-dependent dispersal, where individuals can ‘measure’ local infection prevalence, that is, the relative number of infected hosts. Such a capacity may be a reasonable assumption, if, for example, prevalence is positively correlated with the quantity of a chemical parasite signal present in the environment, or if other signals, such as visual cues can be used to identify infection. Our model allows the free and independent evolution of both components of host dispersal plasticity. Evolutionarily stable levels of plasticity are therefore emergent properties, resulting from epidemiological and evolutionary dynamics in the host-parasite metapopulation. Importantly, our modelling framework takes kin competition into account by default (Poethke et al., 2007). We find that infection-biased dispersal evolves over a wide range of conditions, mainly driven by kin selection, as previously described by Iritani and Iwasa (2014). Going beyond previous theory, we show that the evolution of prevalence-dependent dispersal requires high levels of spatio-temporal autocorrelation in local prevalences, which only arise under high parasite virulence in our model. Under these conditions prevalence-dependent dispersal feeds back on ecological dynamics and stabilizes the host-parasite system.

## The host-parasite model

### Model overview

In this individual-based SI model, infected host individuals (*I*) carry the parasite, and those that are not infected are susceptible (*S*). For simplicity, there is no variation in resistance, and recovery is not possible. The individuals disperse, reproduce and interact in a grid-like landscape (10×10 patches) where each patch is connected to its eight nearest neighbours. In order to avoid edge effects, edges are wrapped around so that the landscape forms a torus. There are no differences in patch quality, and each patch initially has *N*_0_ individuals, half of which are infected implying an initial infection prevalence of 0.5. The host has discrete, non-overlapping generations and reproduces sexually. Dispersal has a genetic basis (Saastamoinen et al., 2018), is natal which implies that individuals may disperse before reproducing, and dispersal may be costly (Bonte et al., 2012b). Parasite transmission is horizontal and parasite dispersal only occurs together with infected hosts. Mating, density regulation, transmission and dispersal decisions are all governed by local rules. Once per generation, patches may suffer form an external local patch extinction. Since we model host-parasite dynamics explicitly, prevalences are not fixed a priori in contrast to Iritani and Iwasa (2014).

We consider two types of dispersal plasticity: First, host dispersal may depend on the infection state (state- or condition-dependence; see Iritani and Iwasa, 2014), that is, the dispersal probability of an individual changes according to whether it is infected or susceptible. Second, the dispersal probability may change according to the local infection prevalence (context-dependence). We therefore assume that the host individuals can estimate local prevalence, as the fraction of infected individuals in their patch without any error (see Bocedi et al., 2012; Poethke et al., 2016a,b, for a treatment of uncertainty during information acquisition). The two types of dispersal plasticity can evolve independently.

### Reproduction and inheritance

The local host population dynamics are modelled explicitly using a discrete-time population growth model, with density-regulation according to Beverton and Holt (1957)

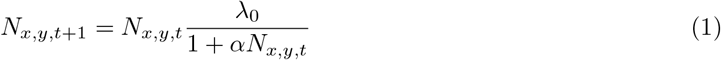

where *λ*_0_ is the fecundity, and *α* is the intraspecific competition coefficient (see Tab. 1 for an overview of parameters and tested values). The equilibrium density in the Beverton-Holt model is given by 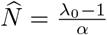.

**Table 1:** Important model parameters, their meaning and tested values.

| Parameter | Description | Values |
| --- | --- | --- |
| $N_0$ | Number of individuals per patch at simulation start | 100 |
| $p_0$ | Initial prevalence per patch | 0.5 |
| $\lambda_0$ | Fecundity | 2, 4 |
| $\alpha$ | Intraspecific competition coefficient. | 0.01 |
| $\mu$ | Dispersal mortality | 0, 0.01, 0.1, 0.5 |
| $m$ | Mutation probability | 0.001 |
| $v$ | Virulence as the reduction in fecundity | 0, 0.1, ..., 1 |
| $\rho$ | Effect of parasite virulence on male fecundity | 1 |
| $\beta$ | Transmission rate | 1, 2, 4 |
| $a$ | Search efficiency of the parasite | 0, 0.001, ..., 0.08 |
| $\epsilon$ | External local patch extinction risk | 0, 0.05 |

After dispersal, the individuals reproduce. Females choose a mate at random from their local patch and produce a number of offspring drawn from a Poisson distribution with a mean 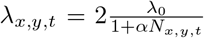 which models demographic stochasticity. The factor of two appears in order to keep *λ*_0_ interpretable at the population level despite half of the population being males which do not produce offspring themselves. In case of a parasite infection *λ_x,y,t_* will be reduced as described below. Each offspring has an equal chance of being either male or female. Since we assume non-overlapping generations, all adults die after reproduction.

An individual’s dependence of dispersal probability on the local prevalence in it’s natal patch (*p_x,y,t_*) and on the individual’s infection state (*X* = *S, I*) is genetically encoded. For convenience, we assume a cubic relationship between dispersal probability and local infection prevalence (technically the four coefficients correspond to four different loci; Eq. 2) that may differ depending on whether individuals are susceptible or infected (state-dependence). This allows the separate evolution of reaction norms for the infected and the susceptible status, and thus represents a simple way to model the simultaneous evolution of infection-dependent and prevalence-dependent evolution.

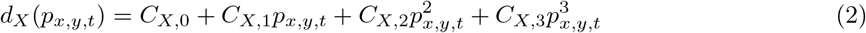

The choice of a cubic reaction norm represents a compromise between a function-valued trait approach that would not make any assumption on the shape of the dispersal reaction norm (Dieckmann and Metz, 2006; Fronhofer et al., 2015) and assuming a functional relationship with fewer parameters. The function-valued trait approach is not feasible here, as preliminary tests showed that the genetic algorithm was not able to optimize efficiently the reaction norm due to the large number of parameters.

Each of the eight modelled loci (four per reaction norm per state) has two alleles, which are inherited independently (that is, we assume no linkage), one from each parent. During inheritance, mutations may occur which is modelled by adding a random number drawn from a normal distribution *N* (0, 0.5) to the parental allelic value with a probability *m* = 0.001, which is the mutation rate. Simulations begin with standing genetic variation present in the population, that is, the coefficients in Eg. 2 assigned to each individual are drawn from a uniform distribution between 0 and 1.

If there is very little variation in local prevalence, for instance, due to the epidemiological dynamics that result from the specific parameter settings of a given simulation, selection on dispersal probability at prevalences that are usually not experienced by the individuals is extremely weak. The reaction norm at these prevalences can therefore not be adequately optimized by our genetic algorithm. Therefore, we only present results of evolutionary stable dispersal reaction norms for the prevalences observed in the respective simulation settings.

### Infection costs and transmission dynamics

Parasites usually utilise host resources for their own growth and reproduction, making their presence costly for the host. In our model, this is captured by the virulence (0 *≤ v ≤* 1) which is a reduction in the mean number of offspring of the host, if infected:

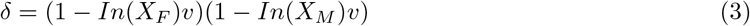

with *In*(*X*) = 0 if *X* = *S* and *In*(*X*) = 1 if *X* = *I*. *X_F_* and *X_M_* are the maternal and paternal infection states, respectively. Therefore, the realised number of offspring is drawn from a Poisson distribution with mean *λ_x,y,t_δ*.

All host individuals are born uninfected and therefore susceptible to infection. The parasite is transmitted after the death of the parental generation. The infection probability for the offspring generation depends on the density of the infected individuals in the parental generation (*I_x,y,t_*), and on the density of all the individuals in the offspring generation (*S_x,y,t_*_+1_). As in Chaianunporn and Hovestadt (2012b), this is represented using a Nicholson-Bailey type model (Nicholson and Bailey, 1935) with a Holling Type II (Holling, 1959b,a) functional response.

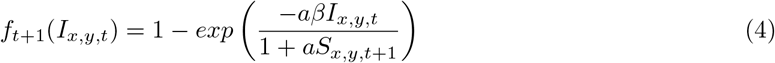

Here, *a* is the search efficiency and *β* is the transmission rate. While originally developed for host-parasitoid systems, search efficiency in our host-parasite model can be seen as the likelihood with which a host individual becomes infected if it encounters a parasite propagule. The transmission rate then governs how many parasite propagules one infected host will release upon its death.

### Dispersal

Dispersal is natal and individuals disperse with a certain probability *d_X_* (*p_x,y,t_*), based on their infection state (*X* = *I, S*) and local prevalence (*p_x,y,t_*) of the disease in the natal patch, calculated according to (2), where each coefficient is the mean of both alleles at a given locus. The target patch is randomly chosen among the eight nearest neighbouring patches. As dispersal is costly (Bonte et al., 2012a), individuals experience mortality during dispersal at a rate *μ*.

### Analysing the effect of kin competition

In an individual-based metapopulation model, kin structure arises by default (Poethke et al., 2007) and is known to influence the evolution of dispersal (Hamilton and May, 1977). To assess the effect of kin selection on the evolution of prevalence- and infection state-dependent dispersal we performed additional simulations using the algorithm described in Poethke et al. (2007), which removes all kin structure from the metapopulation. Briefly, before dispersal and after the transmission of the parasite, all the individuals are shuffled and redistributed across the metapopulation, while maintaining the pre-shuffling density, prevalence, and sex ratio in each patch. This allows all-else-being-equal comparisons of results with and without kin structure.

### Density-dependent dispersal

In general, infection prevalence may be correlated with population density. For instance, high prevalence could imply low host densities due to virulence, and vice versa. Therefore, instead of using prevalence as a source of information to base dispersal decisions on, hosts may evolve to use host population density as information, if population density is less costly to assess. To explore this possibility, we ran additional simulations replacing prevalence-dependent reaction norms by density-dependent reaction norms of dispersal. The dispersal rate was then a function of the local density (*N_x,y,t_*) and infection state (*X* = *S, I*). Poethke and Hovestadt (2002) have shown theoretically that in discrete-time models, such as ours, density-dependent dispersal must follow a threshold function of the form

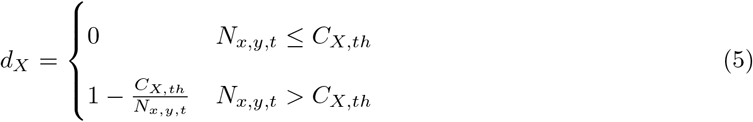

In these simulations the threshold density *C_X,th_* corresponding to each infection state (*X* = *S, I*) can evolve, instead of the polynomial coefficients in Eq. 2. The individuals do not disperse if the local density is below the threshold, and the dispersal probability increases in a saturating way thereafter.

## Results and Discussion

Both infection state- and prevalence-dependent dispersal evolved in our simulations (Fig. 1). Among the different demographic and epidemiological parameters in our model (see Tab. 1 for an overview), parasite virulence was a main factor impacting the evolution of dispersal plasticity qualitatively (see Supplementary Material Fig. S1–S6). For illustration, we have chosen a representative scenario, showing evolutionarily stable levels of dispersal plasticity for low, intermediate and high virulence (*v* = 0.2, 0.5, 0.8, respectively), while other parameters are kept constant (Fig. 1 A–C).

**Figure 1:**
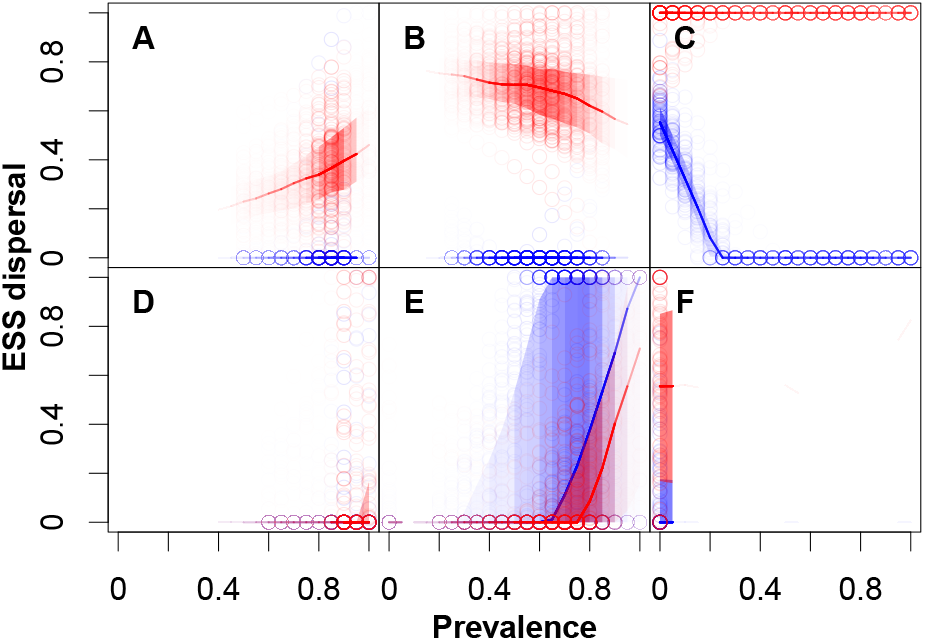
Evolutionarily stable dispersal reaction norms for prevalence- and infection state-dependent dispersal. Red colours represent reaction norms of infected individuals and blue colours show reaction norms when susceptible. The standard and the shuffled (no kin competition) scenarios are represented by the upper (A–C) and lower (D–F) rows, respectively. Virulence increases from left to right. *v* = 0.2 for A and D, *v* = 0.5 for B and E, and *v* = 0.8 for C and F. This graph shows the reaction norms of 100 randomly chosen individuals at the end of the simulation (points) weighted by the normalised frequency of prevalences across space and time, that is, darker shades correspond to higher frequency of prevalence and lighter to lower frequency across space and time. We chose this weighting because the ecological settings of the simulations dictate the prevalences that occur and therefore also whether selection can act on parts of the reaction norm. Our approach therefore guarantees that the results we show are ESS and not the result of drift, for instance, which will dominate for sections of the reaction norm that correspond to prevalences which have never occurred during a given simulation. The lines represent the median dispersal probability corresponding to that prevalence and the shaded regions are the quartiles. Constant model parameters: *λ*_0_ = 4, *α* = 0.01 *β* = 4, *a* = 0.01, *μ* = 0.1, *E* = 0.05.

### Virulence modulates evolution of dispersal plasticity

We find a transition from mainly infection state-dependent dispersal at low to intermediate levels of virulence (*v* = 0.2, 0.5; Fig. 1 A, B) to clearly infection state- and prevalence-dependent dispersal at high virulence (*v* = 0.8; Fig. 1 C). This pattern is not very sensitive to other model parameters, and holds across different values of transmission rate (*β*), parasite search efficiency (*a*), host fecundity (*λ*_0_) and dispersal mortality (*μ*; see also Supplementary Material Fig. S1–S6).

In our example (Fig. 1 A–C) and more generally in all our results, infection state-dependent dispersal always evolves to become *I*-biased, that is, dispersal is induced when individuals become infected. In fact, uninfected individuals practically never disperse, whereas infected dispersal probability is consistently high. The *I*-bias becomes maximal at high virulence, where infection leads to unconditional dispersal (Fig. 1 C). This finding of an *I*-bias confirms the theoretical predictions made by Iritani and Iwasa (2014). We recover these results although, unlike their model, we do not assume a difference in the cost of dispersal for susceptible and infected individuals, nor do we allow for recovery of the infected individuals during dispersal.

Furthermore, Iritani and Iwasa (2014) held prevalences constant across space and time, thus precluding the possibility for the evolution of prevalence-dependent dispersal. By contrast, we implemented the possibility for prevalence dependency of dispersal to evolve. Moreover, infection prevalence was allowed to vary freely from patch to patch, as a consequence of the interaction between dispersal, transmission dynamics (determined by *β* and *a*) and virulence.

In conclusion, a clear pattern of prevalence-dependent dispersal only evolves at high virulence and only for the susceptible state. Uninfected individuals show a steep negative reaction norm, meaning that dispersal is high when prevalence is locally low, but rapidly decreases to zero, as soon as prevalence reaches a threshold (Fig. 1 C). Infected individuals, by contrast, only show moderate trends of plasticity at low and intermediate virulence (Fig. 1 A, B). Below, we will discuss two mechanisms driving the observed evolution of dispersal plasticity.

### Kin competition drives infection state-dependent dispersal

Kin selection can be a major driver of dispersal evolution (Bowler and Benton, 2005), even if dispersal is very costly (Poethke et al., 2007) or in the absence of ecological conditions favouring dispersal (Hamilton and May, 1977). In our model, kin selection provides an intuitive explanation of *I*-biased dispersal: infected individuals leave a patch to reduce effect of kin competition on their uninfected relatives. This is because infected individuals have a lower fitness expectation than susceptible individuals, therefore, the cost of dispersing in terms of inclusive fitness is lower if infected individuals disperse at a higher rate and susceptible individuals do not disperse. Hence, the dispersal rate evolves in such a way that kin competition is reduced (similar to Hamilton and May, 1977) by being infection state-dependent. Indeed, both Poethke et al. (2010) for host-parasitoid and Iritani and Iwasa (2014); Iritani (2015) for host-parasite systems demonstrate that relatedness underlies the evolution of dispersal plasticity in the attacked organism.

In individual-based metapopulation models like ours, kin structure arises by default (Poethke et al., 2007). Its effect on evolutionary outcomes can be tested by using additional simulations with a constantly reshuffled genetic population structure (Poethke et al., 2007). We find that this elimination of kin structure prevents the evolution of a clear dispersal bias, and thus susceptible and infected individuals disperse with similar probabilities for a given prevalence and virulence (Fig. 1 D–F). Unlike Iritani and Iwasa (2014), where *I*-biased dispersal is partly explained by the opportunity for recovery during dispersal, there is no additional direct advantage of dispersal for infected individuals in our model. Thus, given the extreme effect of excluding kin structure, a plausible explanation for our *I*-bias is that the dispersal of infected individuals increases the inclusive fitness of the philopatric susceptible individuals by reducing the effect of kin competition. A priori, this risk should be highest under high virulence, and consequently the evolutionarily stable *I*-bias maximal, which is indeed what we observe in Fig. 1.

One surprising finding is that dispersal bias sometimes evolves at no virulence and a non-zero dispersal mortality, that is, when there is no difference in fecundity between susceptible and infected individuals. Without any external extinction risk (*ϵ* = 0), *S*-biased dispersal evolves for zero and low virulences (Fig. S2). While counter-intuitive at first, we suggest that this is linked to an effect of demographic stochasticity (Travis and Dytham, 1998) acting on susceptible individuals: For high enough search efficiencies, low virulences lead to a high prevalence, which means that the densities of susceptible individuals are greatly reduced. If an individual is susceptible in this scenario, it is more likely to go extinct. This interpretation is supported by simulations with greatly increased equilibrium densities (*λ*_0_ = 4, *α* = 0.001 which implies an equilibrium density of 3000 individuals) where ESS dispersal rates tend towards 0 for both states, infected and non-infected.

### Context-dependent dispersal evolves under conditions of high spatio-temporal variation

Context-dependent dispersal only evolves if there is sufficient spatio-temporal variability in the relevant local condition (Poethke et al., 2016b). More specifically, plasticity is strongly favoured if spatial variation is combined with temporal autocorrelation in local conditions, that is, if conditions are sufficiently predictable for individuals to take the appropriate dispersal decision. We inspected the spatio-temporal variation in infection prevalence, the ‘local context’ in our model, across virulences. At low virulence (*v* = 0.2), prevalence is constantly high and does not show variation between patches (Fig.2), which implies that context-dependent dispersal does not evolve because the context (prevalence) is not variable enough. The range of prevalences broadens for intermediate virulence and is maximal for high virulence, with prevalences ranging from 0 to 1 across the metapopulation (Fig.2). Moreover, only for high virulence, we find strong oscillations in prevalence within patches (Fig.2), leading to generally higher levels of temporal autocorrelation (positive or negative). This combines maximum spatial variation with temporal predictability, both prerequisites for dispersal plasticity to evolve. In other words, ‘knowing’ that low current infection prevalences will predictably change to high levels, explains the evolution of increased dispersal of susceptible individuals at low prevalence (Fig. 1 C), because this decision can be expected to yield an inclusive fitness benefit. Conversely, the rule not to disperse at high local prevalences means that staying in the local patch and waiting for better conditions (i.e., lower prevalences) is more rewarding than leaving the patch.

We observe that when there is no external extinction risk (*ϵ* = 0) and low parasite search efficiency (*a* = 0.005), even high virulences (*v* = 0.8) may not lead to the evolution of prevalence-dependent dispersal, even though the entire range of prevalences occur (Fig. S2). Under these conditions, we observe a positive temporal autocorrelation in prevalences within a patch (see Fig. S8). Therefore, susceptible individuals stay in their natal patch at low prevalences because it is likely to remain that way giving them an inclusive fitness advantage. The spatial variation in prevalences implies that if individuals disperse, they are more likely to reach a patch with a higher prevalence, causing a greater infection risk for their offspring which reduces their fitness expectation (Hastings, 1983). As the ESS emigration rate depends on the availability of good habitat in the metapopulation (Poethke et al., 2011) this selects against dispersal at low prevalence.

### Evolution of infection state- and density-dependent dispersal

Our model uses infection prevalence for context-dependent dispersal. However this information may not always be easily obtained by individuals. Population density is another, potentially more accessible cue that can drive context-dependent dispersal (Travis and Dytham, 1999; Poethke and Hovestadt, 2002; Matthysen, 2005; Nowicki and Vrabec, 2011). Because host density is also likely to be sensitive to parasite infestation, we re-ran our model using the density-dependent dispersal function derived by Poethke and Hovestadt (2002) as a reaction norm keeping the possibility for the evolution of state-dependency of dispersal. As above, we find the evolution of a consistent *I*-bias of dispersal (Fig. 3 A–C), showing that state-dependent dispersal evolves independently of the type of context-dependency.

We also find density-dependent dispersal of susceptible individuals under high virulence. But unlike for prevalence dependency (Fig. 2 C), the relationship is positive: dispersal increases with increasing population density (Fig. 3 C). These mirrored patterns can be explained by the fact that density and prevalence are negatively correlated (Fig. 3 D–F): host populations become more decimated at higher infection prevalence if virulence is sufficiently high. Obviously, just like for prevalence, density needs to fluctuate in a predictable way for plasticity to evolve, which only occurs in the high-virulence scenario (Fig. 3 F). This finding implies that hosts may use either density or prevalence as a cue for context-dependent dispersal, depending on which information is less costly to obtain or more reliably estimated.

**Figure 2:**
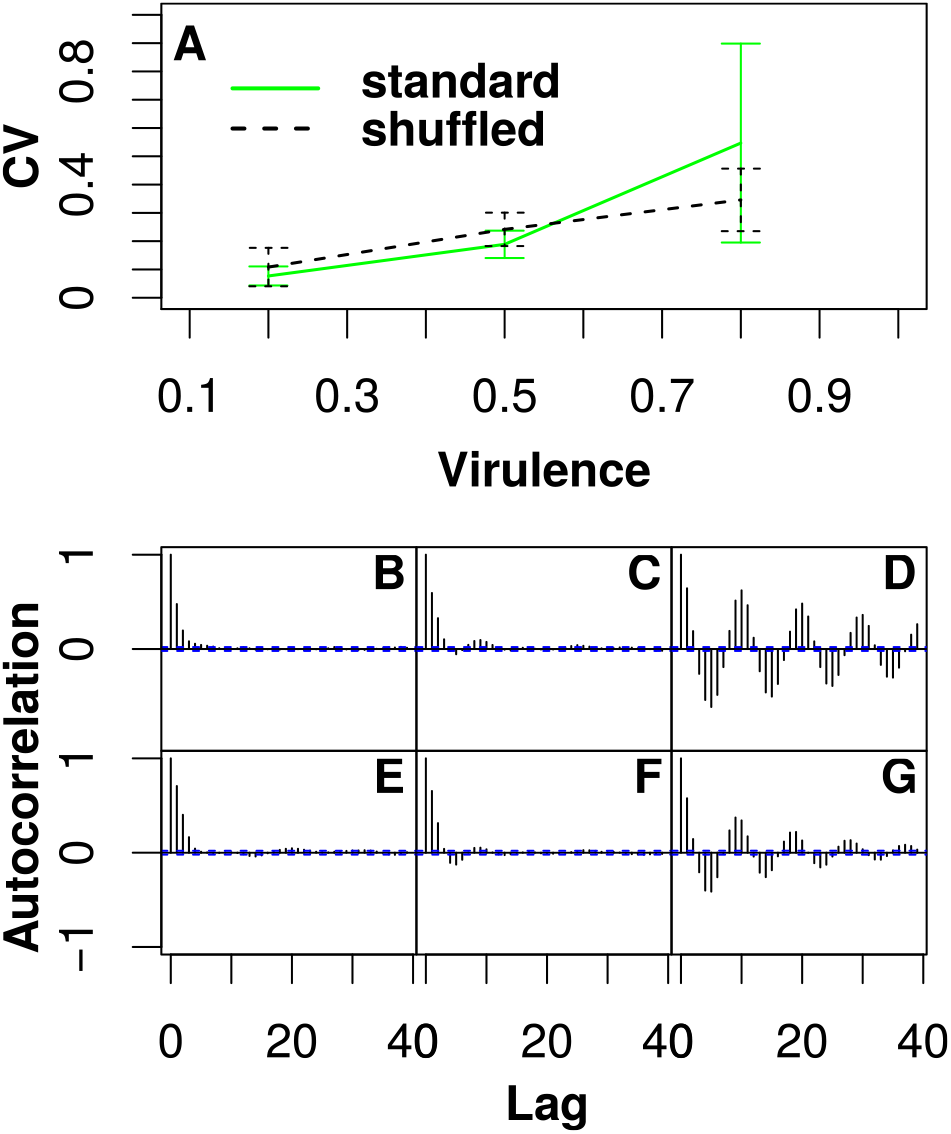
Spatio-temporal variation in local conditions, that is, prevalence. A shows the spatial coefficient of variation (CV) of the prevalence with increasing virulence. The standard scenario is represented by green colour and solid lines and the shuffled scenario (no kin competition) is shown as black dashed lines. Error bars represent standard deviation of the CV across 10 simulation replicates. B–G visualise the temporal autocorrelation coefficient as a function of the time lag for one example patch, with B– D and E–G as the standard and shuffled scenarios, respectively. Virulence changes from left to right: *v* = 0.2, 0.5, 0.8. Constant model parameters: *λ*_0_ = 4, *α* = 0.01 *β* = 4, *a* = 0.01, *μ* = 0.1, *ϵ* = 0.05.

**Figure 3:**
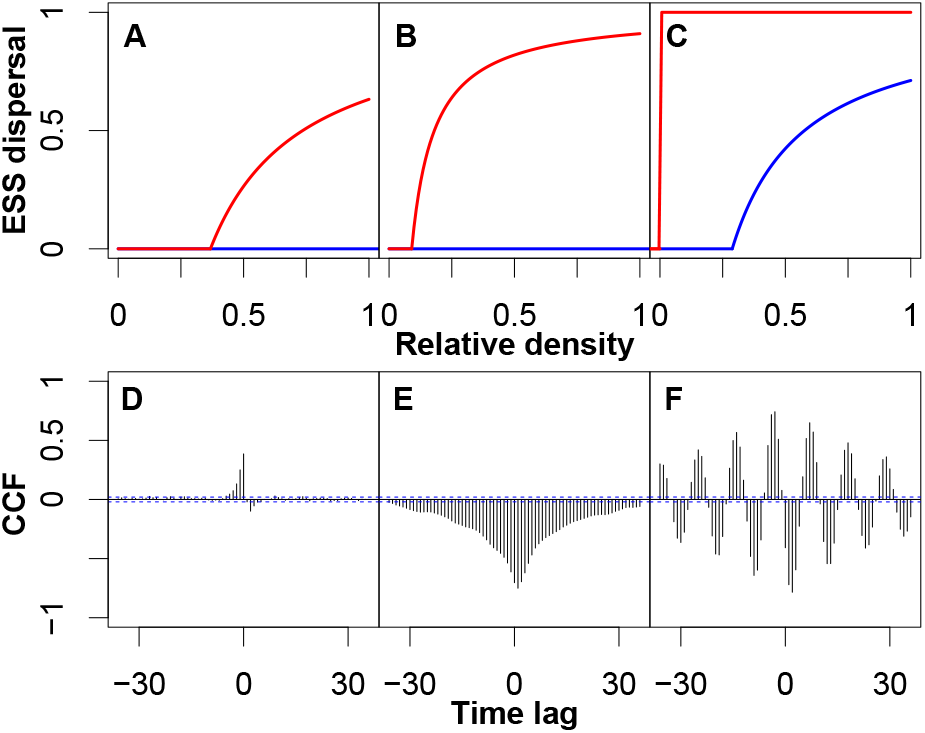
Evolutionarily stable dispersal reaction norms for density- and infection state-dependent dispersal (A–C). Red colours represent reaction norms of infected individuals and blue colours show reaction norms when susceptible. The x-axis indicates population density relative to the equilibrium population density. The lower row (D–F) shows the cross correlation between host population density and prevalence. Virulence increases from left to right (D: *v* = 0.2 E: *v* = 0.5, F: *v* = 0.8). Constant model parameters: *λ*_0_ = 4, *α* = 0.01 *β* = 4, *a* = 0.01, *μ* = 0.1, *ϵ* = 0.05.

Even though density and prevalence are generally negatively correlated, the relationship may not always be that simple. Due to the dynamic nature of population regulation by the parasite, prevalence and density are not perfectly negatively correlated, such that high population density may at times be associated with high prevalence, but at others with low prevalence. We show here that distinct patterns of cross correlations may arise, that is, prevalence and density are correlated with a time lag (Fig. 3 D–F).

### Eco-evolutionary feedbacks

Context-dependent dispersal based on density or on prevalence cues may have different demographic consequences at the metapopulation level, that is, eco-evolutionary feedbacks. For instance, at ESS, patch extinction frequency, population density and prevalence across time can be impacted differentially (Fig. 4).

**Figure 4:**
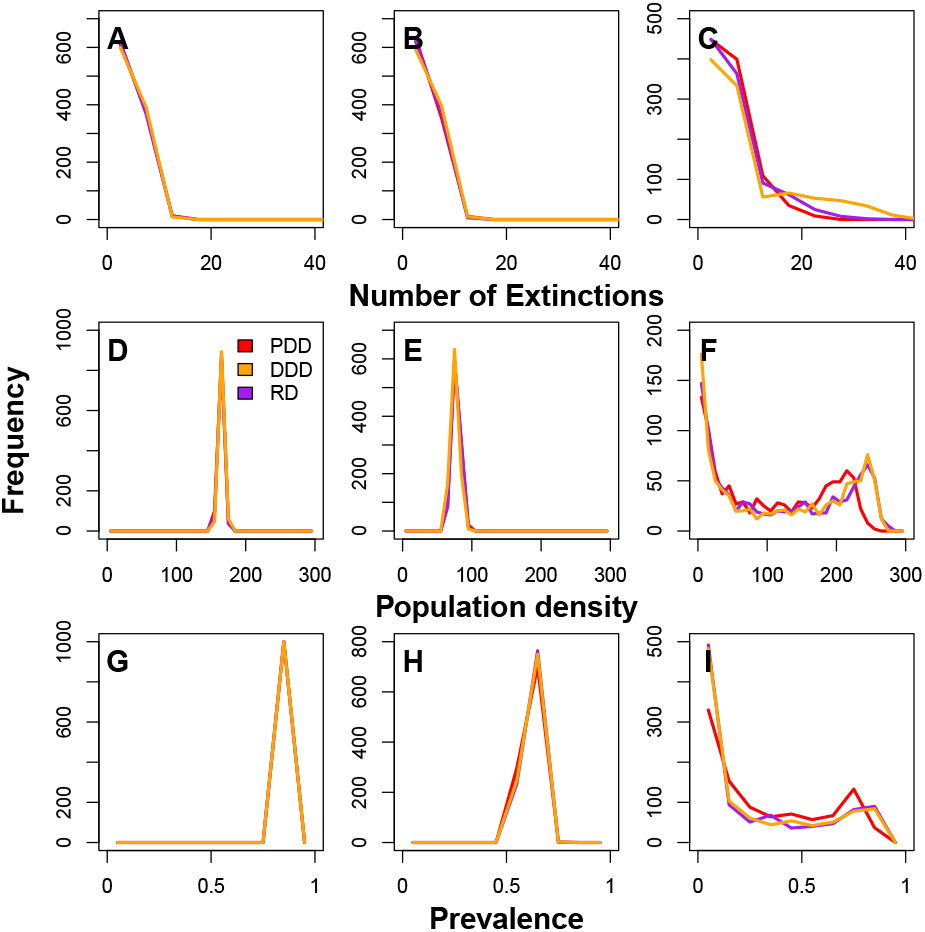
The distribution (frequency) of number of extinct patches (due to external local patch extinctions *E* and emergent from the host-parasite dynamics), median prevalences and host densities at a given time for prevalence-dependent dispersal (PDD; red), density-dependent dispersal (DDD; orange) and the ecological scenario (RD; purple), for the last 1000 time steps. The ecological scenario is shown as a comparison and assumes that the dispersal rate is constant but equal to the observed overall ESS dispersal rate in the prevalence-dependent case. *v* = 0.2, 0.5, 0.8 from left to right for the focal scenario (*λ*_0_ = 4, *β* = 4, *μ* = 0.1, and *ϵ* = 0.05, *a* = 0.01). As expected, the results for *v* = 0.2 and *v* = 0.5 are very similar between the dispersal scenarios as under these conditions context-dependent dispersal does not evolve. For *v* = 0.8, the main effect of prevalence-dependent dispersal is a stabilizing effect: prevalence-dependent dispersal reduces the number of extinctions as well as extremes in host density and in prevalence.

Under the high-virulence scenario, where we observe the evolution of prevalence-dependent dispersal (Fig. 1 C), prevalence-dependency feeds back on demography and leads to a lower overall patch extinction rate than density-dependent dispersal, and to a higher frequency of intermediate prevalences at equilibrium (Fig. 4 and Figs. S9 – S10). Therefore, while not reducing overall mean levels of prevalence, infection state- and prevalence-dependent dispersal reduces the occurrence of extremes, both in terms of prevalences and densities (Fig. 4). Prevalence-dependent dispersal is therefore stabilizing and may even be seen as a “spatial tolerance” mechanism which can be relevant when the evolution of resistance is not possible. Investigating the concurrent evolution of resistance and other relevant life-history traits is of course an important next step. In summary, the demographic conditions defined by the host-parasite interaction drive the evolution of context-dependent dispersal which then feeds back on these demographic conditions in an eco-evolutionary feedback loop.

## Conclusion

In conclusion, we show that *I*-biased, state-dependent dispersal readily evolves in our model and is pervasive. The dispersal bias evolves due to kin selection which is consistent with previous studies (Iritani and Iwasa, 2014; Iritani, 2015).

Interestingly, empirical studies often show the opposite pattern (*S*-bias; Cameron et al., 1993; Bradley and Altizer, 2005; Nørgaard et al., 2019), which may be explained by infections weakening hosts and thereby reducing their dispersal capacity. Alternatively, Iritani and Iwasa (2014) show that *S*-bias may evolve under specific conditions such as strong virulence affecting competitive ability, high rates of parasite release during dispersal, or low virulence for infected emigrants. While we did not include these specificities in our study for reasons of simplicity, our model can be extended and applied to understand dispersal bias and context-dependent dispersal in more complex host-parasite systems, also including vertical transmission and different modes of sexual reproduction, for example. In contrast to previous work, we relax the strong assumption made for instance by Iritani and Iwasa (2014) of constant global prevalence, no transmission parameters and fixed population dynamics. Importantly, our approach allows for eco-evolutionary feedbacks (Govaert et al., 2019) to occur in our model (Fig. 4).

In contrast to state-dependent dispersal, we find rather restricted conditions for prevalence-, that is, context-dependent dispersal to evolve. Prevalence-dependent dispersal requires particular life-history trait values, such as high virulence, and a sufficient degree of predictability of epidemiological dynamics to evolve. In addition, the observed evolutionary outcome, a negatively prevalence-dependent dispersal reaction norm if susceptible, seems counter-intuitive at first. One may have naively predicted that uninfected individuals should only leave a patch if local prevalence is too high. In predator-prey systems, this seems to be a general pattern as recently demonstrated by Fronhofer et al. (2018). However, even in predator-prey systems, theory shows that the slope of the reaction norm will depend on the spatial and temporal autocorrelation of predation risk (Poethke et al., 2010) as in our present study. Importantly, in our eco-evolutionary model, host-parasite dynamics and spatio-temporal heterogeneity in prevalences are a result of transmission which depends on the density of infected and susceptible individuals as well as on virulence, along with feedbacks from evolved dispersal strategies. The emergent spatial variation in prevalences, temporal autocorrelation and oscillation in density drive the evolution of prevalence-dependent dispersal. Vice versa, the evolutionarily stable dispersal strategy modifies ecological dynamics. Specifically, prevalence-dependent dispersal has a stabilizing effect (Fig. 4; see also Fronhofer et al. 2018 for similar findings in food chains).

Finally, we show that context-dependency in dispersal may evolve even if information on prevalences is not available, too costly to acquire or too noisy. Due to cross-correlations between prevalence and density, individuals may use local population density as an alternative source of information for context-dependent dispersal in host-parasite systems.

## Author contributions

J.N.D., O.K. and E.A.F. conceived the study. J.N.D. and E.A.F. developed the models. J.N.D. analysed the models. J.N.D. wrote the manuscript and all authors commented on the draft.

## Acknowledgements

This is publication ISEM-YYYY-XXX of the Institut des Sciences de l’Evolution – Montpellier.

## Data availability

Simulation code will be made available via GitHub and a Zenodo DOI.

## Supplementary Material

**Figure S1:**
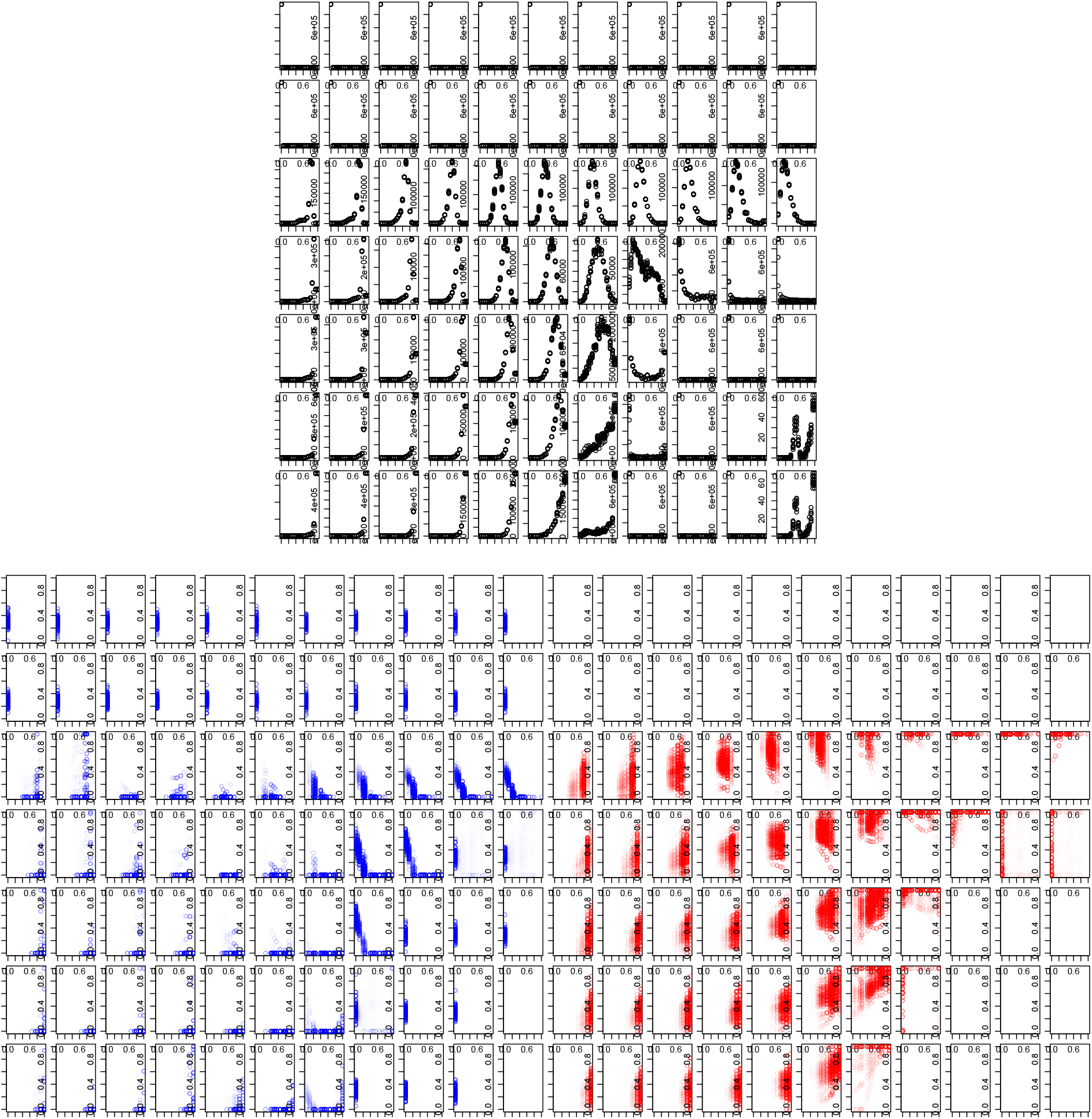
Focal Scenario (*λ*_0_ = 4, *β* = 4, *μ* = 0.1, and *ε* = 0.05). Top: Histograms of local prevalences of all patches, over all time. Down: Left-Dispersal probability as a function of prevalence, if susceptible, right-if infected. Global x-axis: *v* = 0, 0.1, *…*, 1 from left to right, Global y-axis: *a* = 0, 0.001, 0.005, 0.01, 0.05, 0.08 from top to bottom

**Figure S2:**
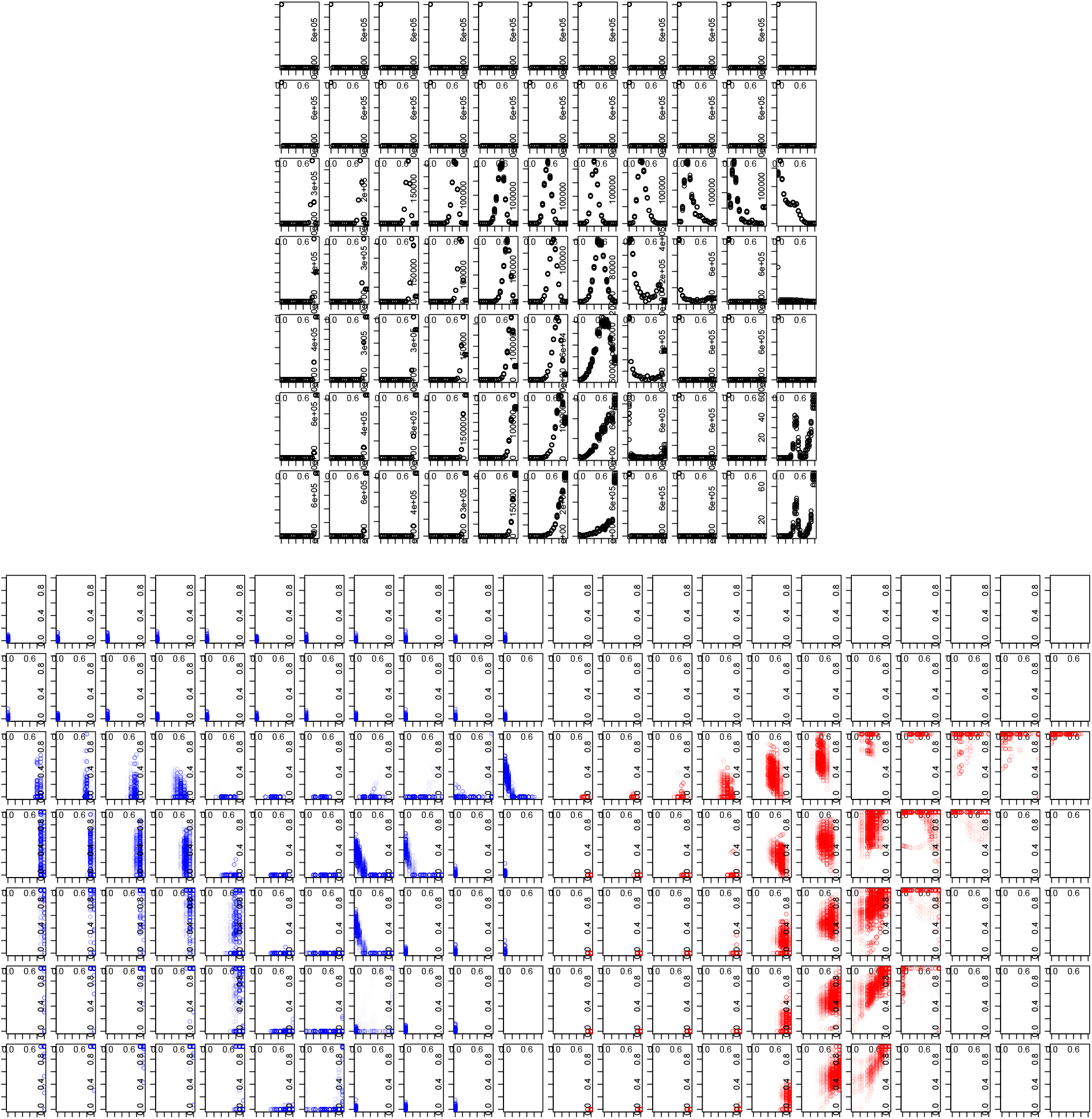
Effect of Extinction risk. (*λ*_0_ = 4, *β* = 4, *μ* = 0.1, and *ε* = 0). Top: Histograms of local prevalences of all patches, over all time. Down: Left-Dispersal probability as a function of prevalence, if susceptible, right-if infected. Global x-axis: *v* = 0, 0.1, *…*, 1 from left to right, Global y-axis: *a* = 0, 0.001, 0.005, 0.01, 0.05, 0.08 from top to bottom

**Figure S3:**
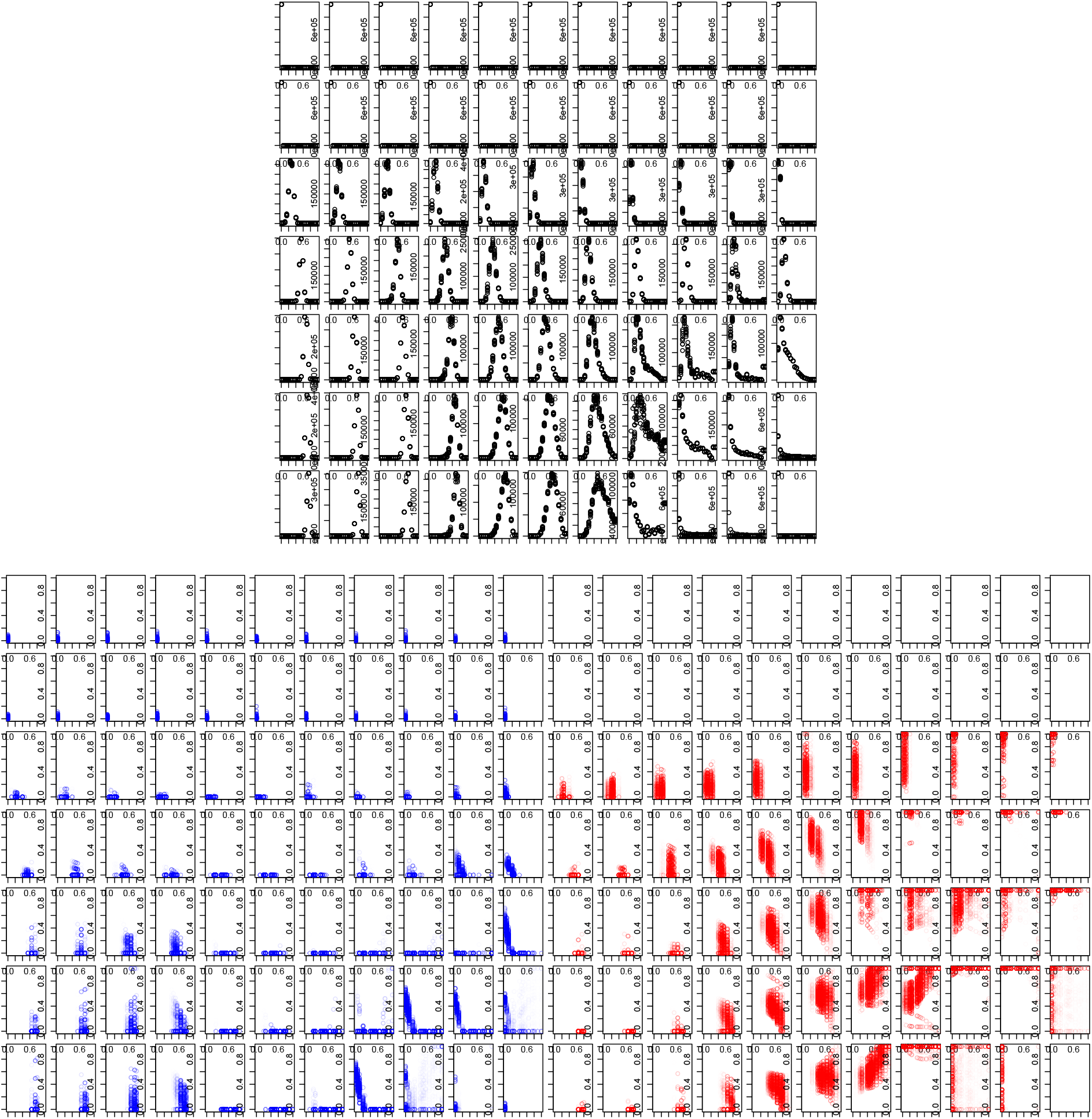
Effect of Transmission rate. (*λ*_0_ = 4, *β* = 2, *μ* = 0.1, and *ε* = 0). Top: Histograms of local prevalences of all patches, over all time. Down: Left-Dispersal probability as a function of prevalence, if susceptible, right-if infected. Global x-axis: *v* = 0, 0.1, *…*, 1 from left to right, Global y-axis: *a* = 0, 0.001, 0.005, 0.01, 0.05, 0.08 from top to bottom

**Figure S4:**
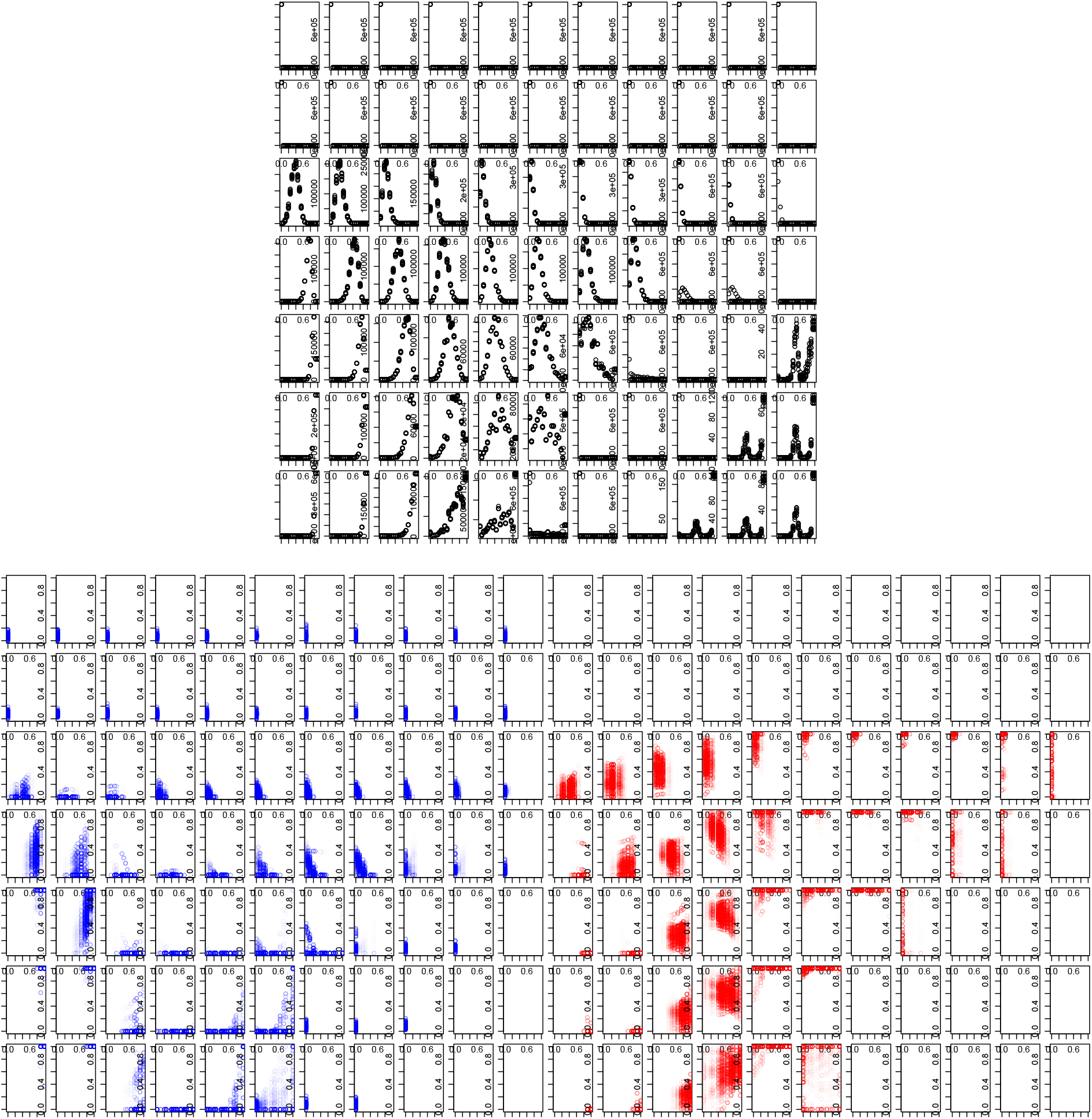
Effect of fecundity. (*λ*_0_ = 2, *β* = 4, *μ* = 0.1, and *ε* = 0). Top: Histograms of local prevalences of all patches, over all time. Down: Left-Dispersal probability as a function of prevalence, if susceptible, right-if infected. Global x-axis: *v* = 0, 0.1, *…*, 1 from left to right, Global y-axis: *a* = 0, 0.001, 0.005, 0.01, 0.05, 0.08 from top to bottom

**Figure S5:**
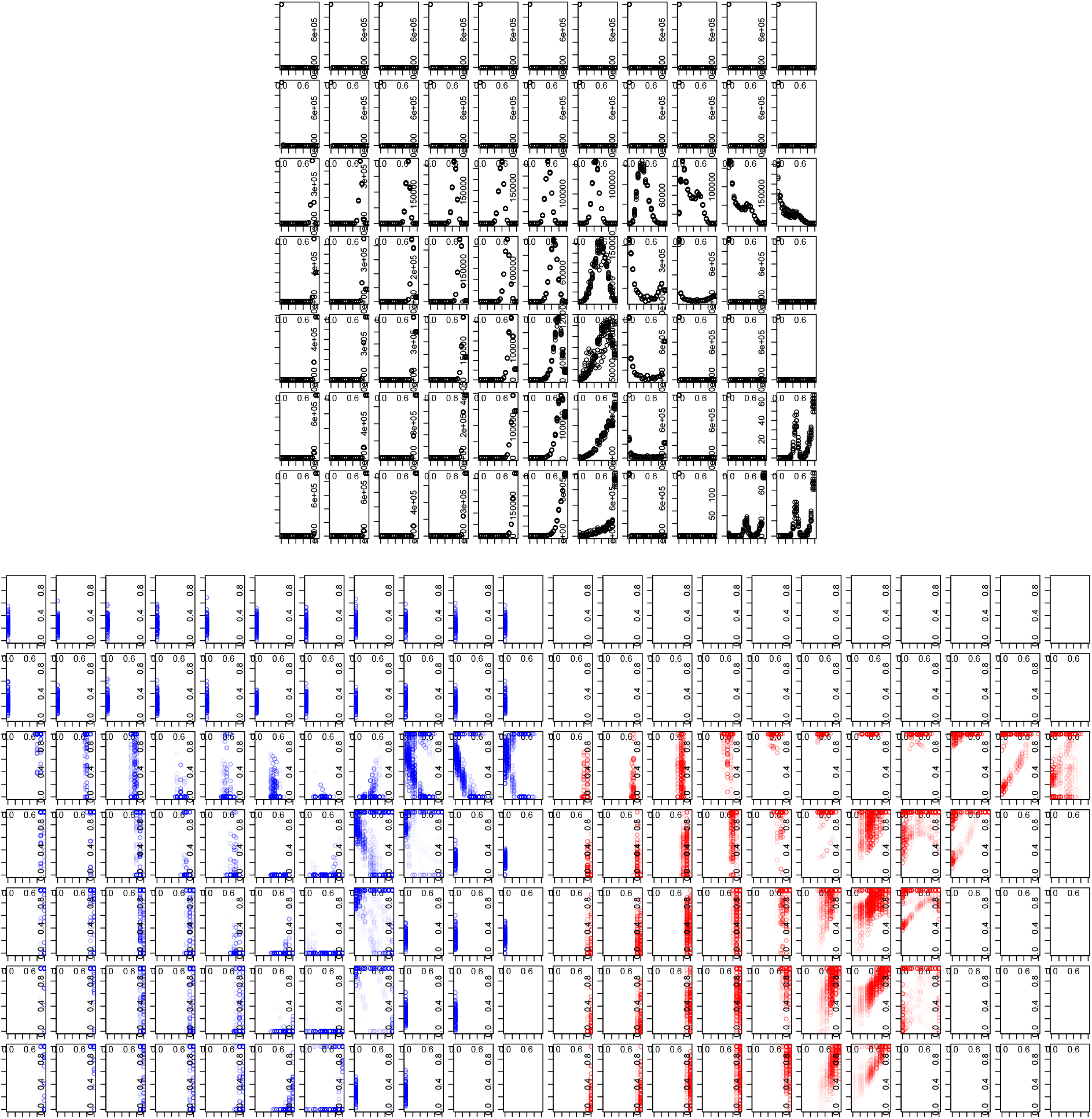
For lower dispersal mortality. (*λ*_0_ = 4, *β* = 4, *μ* = 0.01, and *ε* = 0). Top: Histograms of local prevalences of all patches, over all time. Down: Left-Dispersal probability as a function of prevalence, if susceptible, right-if infected. Global x-axis: *v* = 0, 0.1, *…*, 1 from left to right, Global y-axis: *a* = 0, 0.001, 0.005, 0.01, 0.05, 0.08 from top to bottom

**Figure S6:**
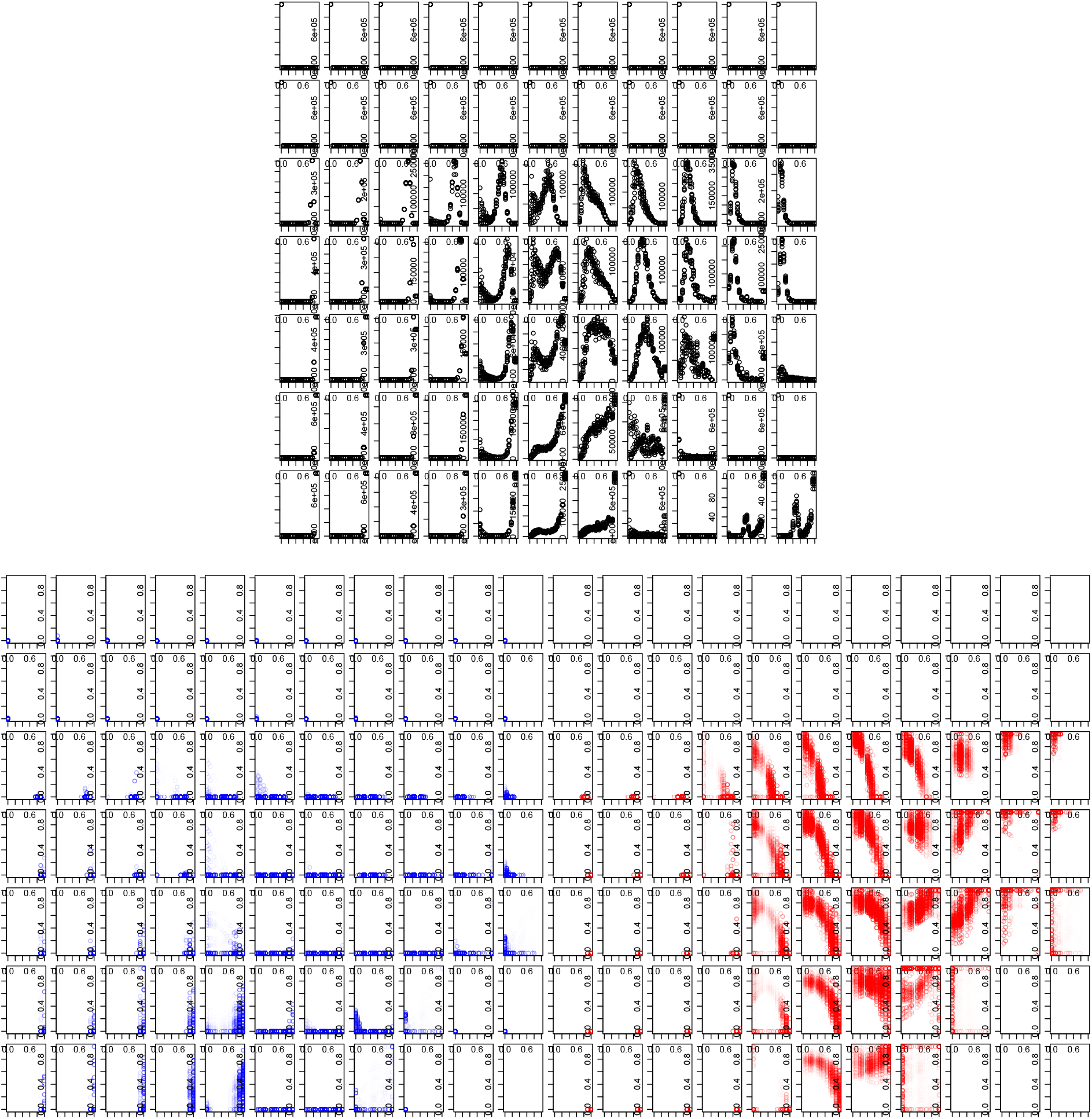
For a higher dispersal mortality (*λ*_0_ = 4, *β* = 4, *μ* = 0.5, and *ε* = 0). Top: Histograms of local prevalences of all patches, over all time. Down: Left-Dispersal probability as a function of prevalence, if susceptible, right-if infected. Global x-axis: *v* = 0, 0.1, *…*, 1 from left to right, Global y-axis: *a* = 0, 0.001, 0.005, 0.01, 0.05, 0.08 from top to bottom

**Figure S7:**
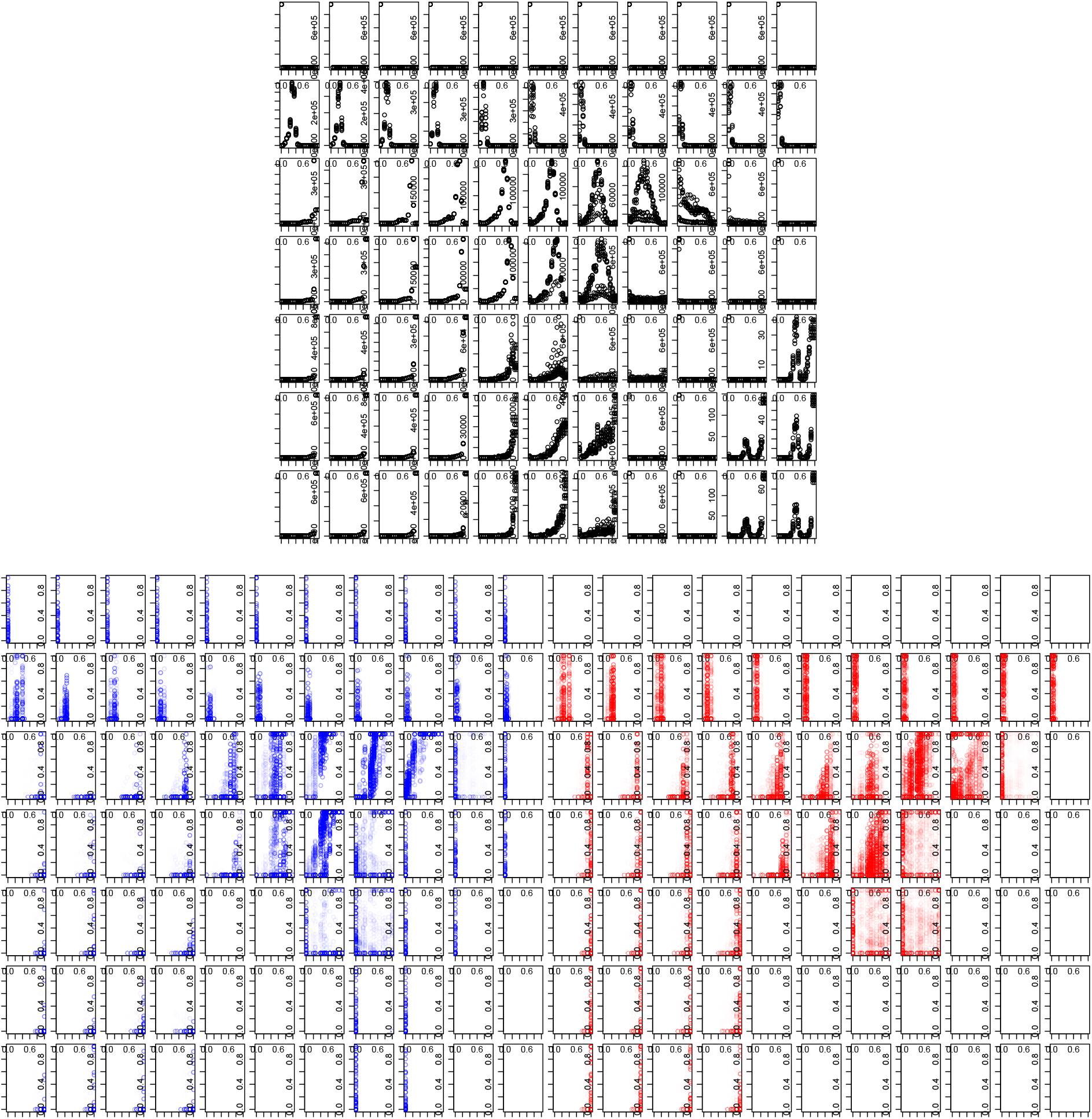
Focal Scenario without kin competition (*λ*_0_ = 4, *β* = 4, *μ* = 0.1, and *ε* = 0.05). Top: Histograms of local prevalences of all patches, over all time. Down: Left-Dispersal probability as a function of prevalence, if susceptible, right-if infected. Global x-axis: *v* = 0, 0.1, *…*, 1 from left to right, Global y-axis: *a* = 0, 0.001, 0.005, 0.01, 0.05, 0.08 from top to bottom

**Figure S8:**
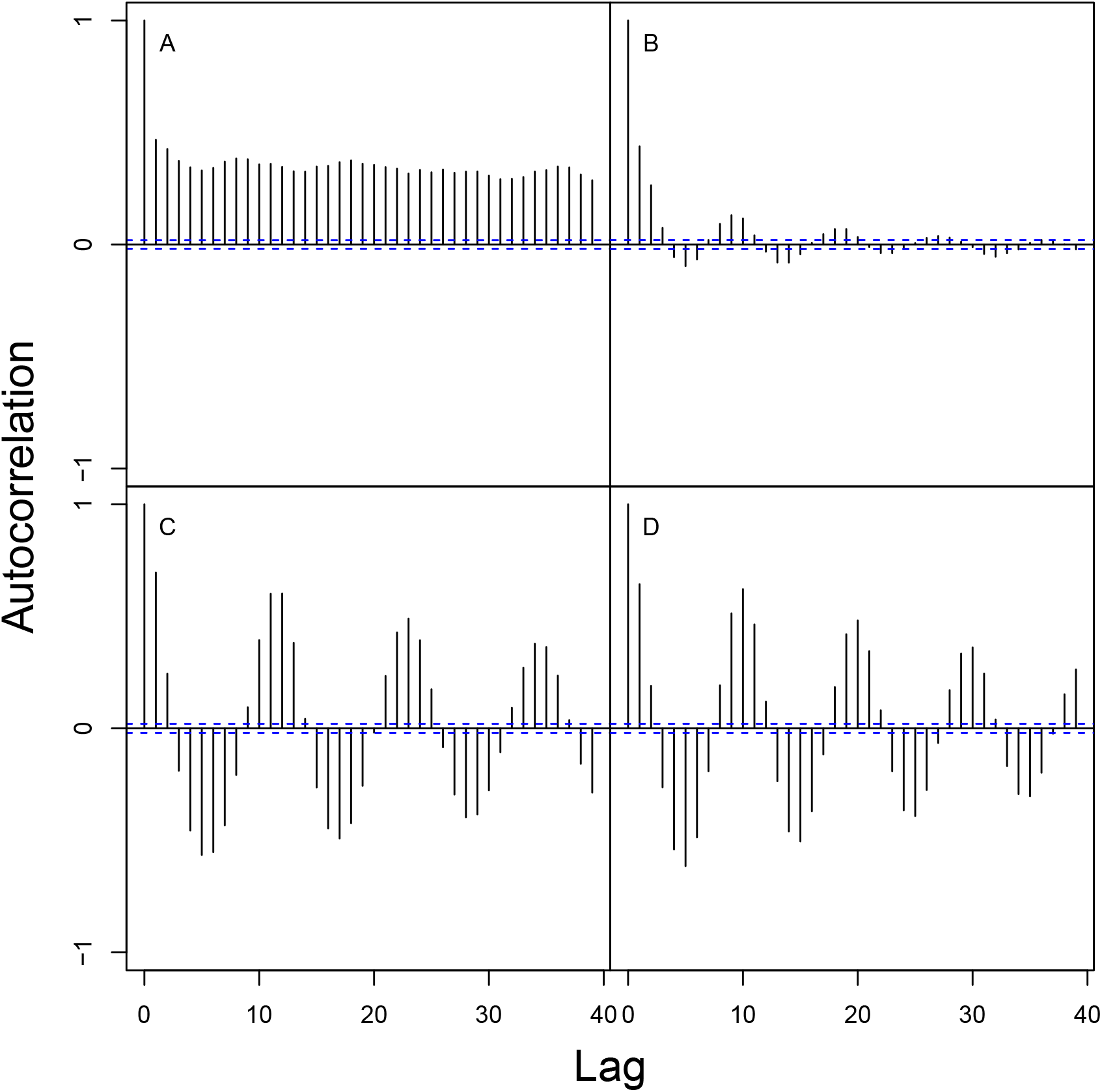
Autocorrelation of prevalences for the high virulence case, for different search efficiency (*a* = 0.005, 0.01 from left to right, and external extinction risk *ε* = 0, 0.05). *λ*_0_ = 4, *β* = 4, *μ* = 0.1, *v* = 0.8). Context-dependency evolves in the cases where there are oscillations (**B,C,D**), and does not evolve when the oscillations are absent (**A**).

**Figure S9:**
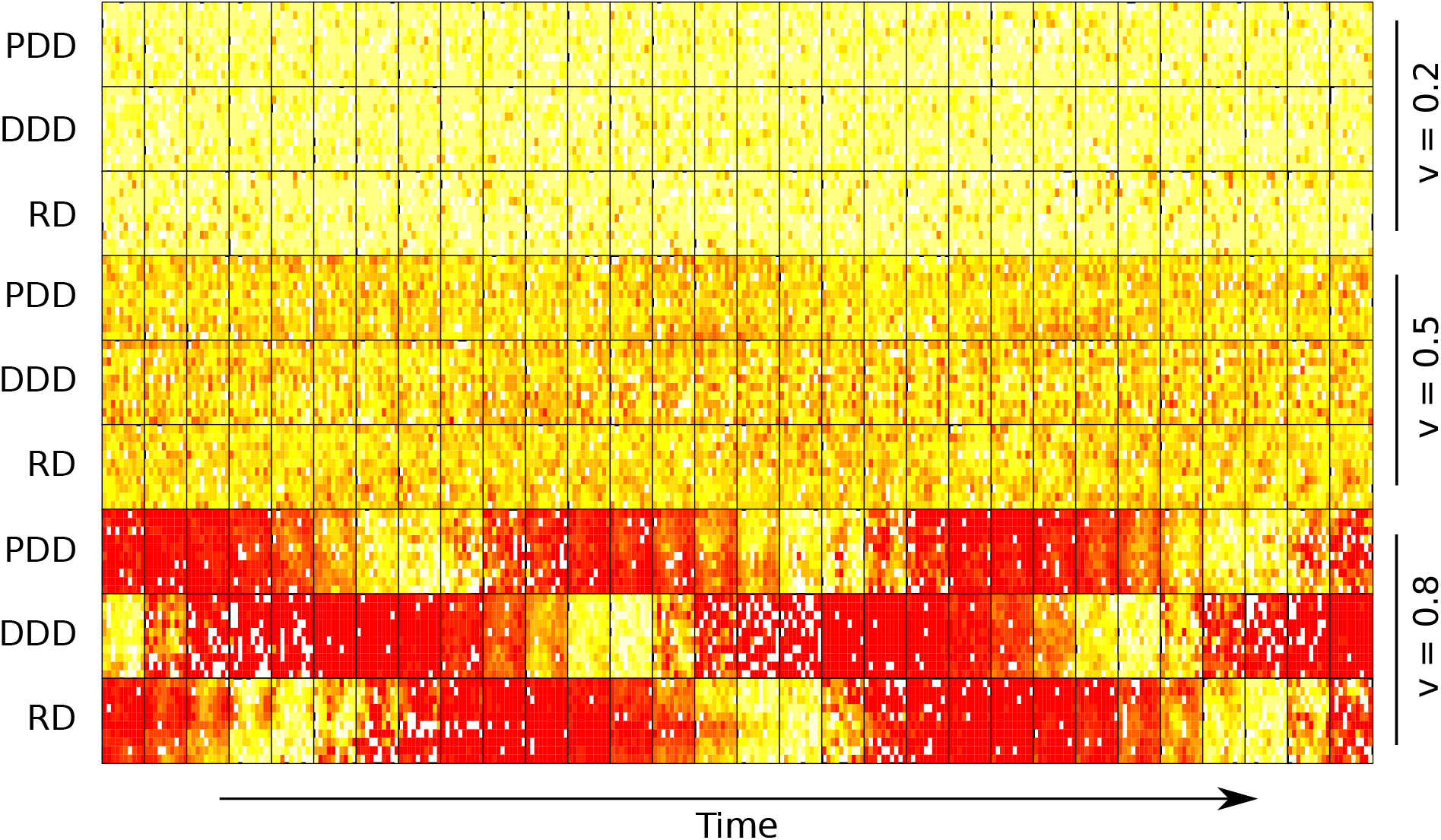
This figure shows the prevalence in each landscape for the last thirty time steps(from left to right). From top to down: the prevalence-dependent (PDD), density-dependent (DDD) and ecological scenarios (RD) are represented one after the other, for *v* = 0.2, 0.5, 0.8 for the focal scenario (*λ*_0_ = 4, *β* = 4, *μ* = 0.1, and *ε* = 0.05, *a* = 0.01).

**Figure S10:**
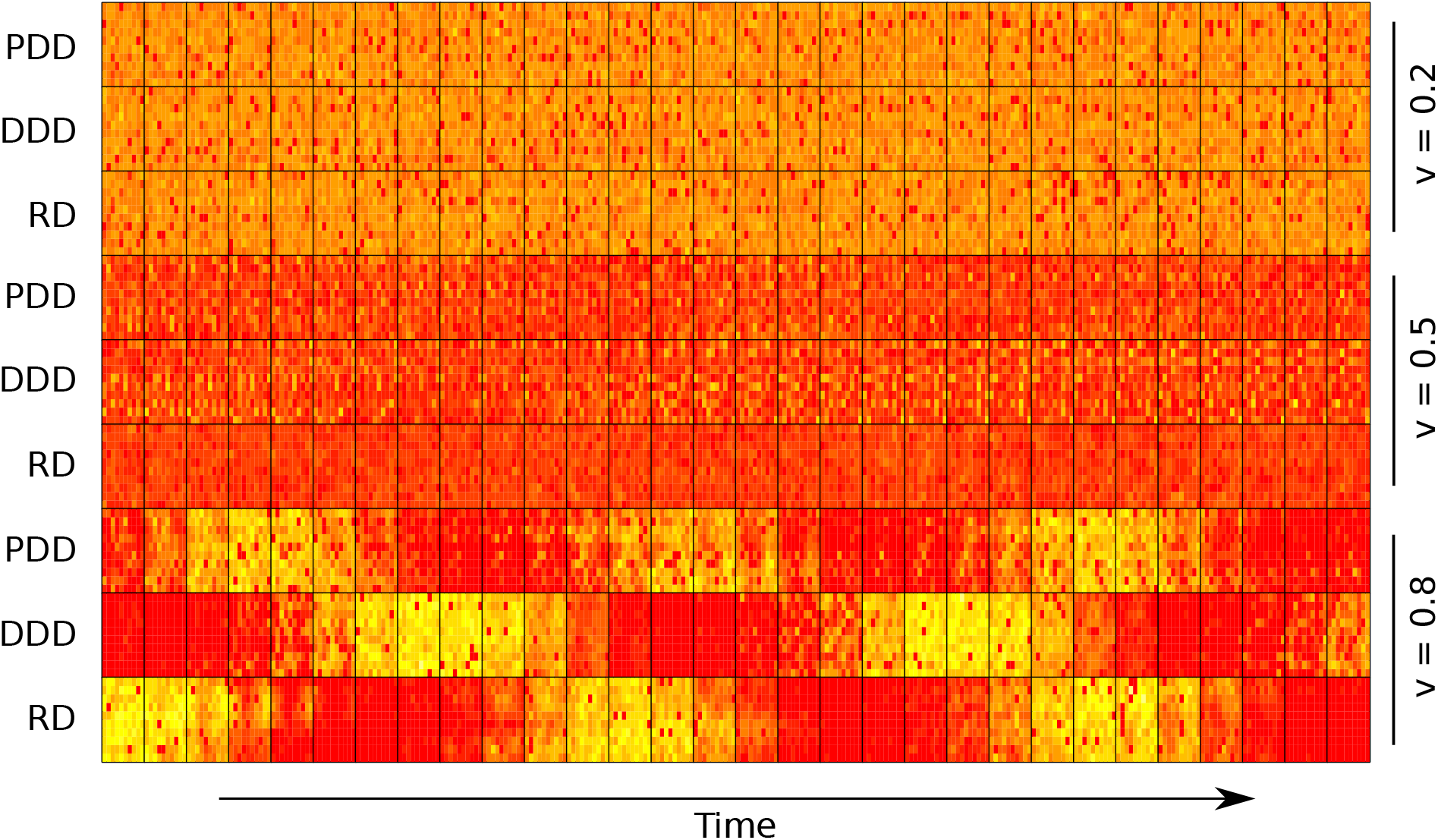
This figure shows the density in each landscape for the last thirty time steps(from left to right). From top to down: the prevalence-dependent (PDD), density-dependent (DDD) and ecological scenarios (RD) are represented one after the other, for *v* = 0.2, 0.5, 0.8, for the focal scenario (*λ*_0_ = 4, *β* = 4, *μ* = 0.1, and *ε* = 0.05, *a* = 0.01).

